# Language Dysfunction Associated with Pathological Brain Connectivity in Children with Epilepsy

**DOI:** 10.64898/2026.07.26.740747

**Authors:** Xiwei She, Olivia Peony, Wendy Qi, Miguel Menchaca, Kerry C. Nix, Wei Wu, Zihuai He, Fiona M. Baumer

**Author notes:** Corresponding Author: Fiona M. Baumer, 290 Campus Drive, E259, Stanford, CA 94305.

## Abstract

Language impairment is common in childhood epilepsy and may arise in part from disruption of the distributed brain networks that support language. In self-limited epilepsy with centrotemporal spikes (SeLECTS)—the most common focal epilepsy of childhood—hyperconnectivity has been linked to poor language outcomes, but the specific network patterns associated with language dysfunction remain unclear, limiting the ability to target neurostimulation rationally.

We recorded high-density EEG from 27 children with SeLECTS and 29 age-matched controls during verb generation and rest, and quantified functional connectivity across multiple frequency bands and bilateral frontal, temporal, occipital, and motor regions. Using multivariate pattern analysis, we identified connectivity patterns that predicted language ability, distinguished patterns shared across groups from those specific to SeLECTS and tested whether spatially specific connectivity provided information beyond whole-brain or hemispheric averages and conventional clinical variables.

Frontal and occipitotemporal connectivity, particularly within the left hemisphere, predicted language ability across groups, whereas motor-network connectivity emerged as the dominant SeLECTS-specific predictor, linking the epileptogenic network to language dysfunction. Connectivity between specific regions outperformed averaged connectivity measures and predicted language beyond epilepsy diagnosis and antiseizure medication use. Task-based connectivity also outperformed resting-state connectivity. These findings show that language ability is associated with distributed yet spatially specific patterns of brain connectivity, while epilepsy introduces distinct alterations centered on the epileptogenic network. Identifying these disease-specific network patterns provides mechanistic insight into language dysfunction and a rational basis for spatially targeted neuromodulation.

## 1. Introduction

Language problems reduce quality of life for millions of children with epilepsy^1,2^. Causes of language impairment in epilepsy are numerous, and disrupted connectivity between cortex critical for language processing is an interesting etiology as it may be amenable to treatment^3–5^. Children with epilepsy frequently exhibit altered functional connectivity compared to typically developing peers^6–8^, with many but not all studies linking elevated connectivity to worse language performance^9–11^. While non-invasive neurostimulation can modulate functional connectivity^12^, stimulation therapies have not been developed to target connectivity-related network disruption and improve language outcomes. A critical and unanswered question is *which* connectivity patterns are specifically linked to language impairment and *how* neurological disease alters these relationships. We hypothesize that children with epilepsy share some language-relevant connectivity patterns with typically developing children, reflecting general principles of language network organization, while also exhibiting epilepsy-specific patterns that may represent promising neurostimulation targets.

Testing this hypothesis requires methods capable of overcoming several challenges. First, language is a complex and distributed process, yet prior work has relied predominantly on univariate analyses that evaluate one connection at a time and cannot capture the distributed structure of language networks^13^. Second, to be clinically useful, connectivity must contribute unique information about language not captured by clinical variables alone. Children with epilepsy generally have worse language performance, particularly those taking antiseizure medications (ASMs)^3,14^, and these same clinical factors are associated with altered connectivity^15–17^. To be a useful treatment target, connectivity must not simply be a proxy for diagnosis or ASM status. Third, connectivity must provide spatially specific information. Prior EEG studies in children with focal epilepsy have reported elevated connectivity globally^6,10,18,19^. If connectivity between specific regions is more important than global averages, this would implicate focal network disruption that could inform spatially targeted intervention. Multivariate pattern analysis (MVPA) addresses all three of these challenges by leveraging machine learning to identify predictive patterns across many connectivity features simultaneously^20,21^.

Here, we determined if brain connectivity, as measured by EEG from children with and without epilepsy performing both verb generation and resting tasks, predicts language ability. We recruited children with self-limited epilepsy with centrotemporal spikes (SeLECTS), the most common focal epilepsy of childhood^22^, because seizures are infrequent and children are typically otherwise healthy and yet have mild-to-moderate language difficulties^23^. Children with SeLECTS have interictal epileptiform discharges (IEDs) originating in the sensorimotor cortices that are associated with both hyperconnectivity and language dysfunction^24^. Connectivity was measured as the weighted phase lag index (wPLI)^25^ across four frequency bands and ten regions of interest spanning superior frontal, interior frontal, temporal, occipital, and motor areas across both hemispheres. These regions were selected to sample the brain widely while capturing core nodes in the language network and motor regions implicated both in SeLECTS pathophysiology and in phonological processing^26–28^. Phonological awareness, a core component of language development that is consistently impaired in children with SeLECTS^14^, was our primary language ability measure. We designed our primary MVPA model to include both connectivity features shared across groups and epilepsy-specific interaction terms. We also tested whether connectivity predicts language beyond clinical variables and whether connectivity-language associations are spatially specific.

## 2. Methods

### 2.1 Participants

Right-handed, 5–13 year-old children were recruited from Lucile Packard Children’s Hospital or the surrounding community. Children with SeLECTS^22^ were identified based on appropriate seizure semiology (hypersalivation, facial/hemibody twitching, or nocturnal tonic-clonic) and centrotemporal IEDs on EEG. Age-matched typically developing controls had no history of seizures or ASM use. Exclusion criteria for both groups included a history of prematurity (<35 weeks); severe neurological or systemic conditions (e.g., stroke, traumatic brain injury, cardiac, or oncologic disease), or focal deficits on neurologic exam. This study was approved by the Stanford University Institutional Review Board, with written consent obtained from parents and assent from children over 7 years old.

### 2.2 Neuropsychological Assessment

Participants completed subtests required for the Phonological Awareness Composite score from the Comprehensive Test of Phonological Processing, Second Edition (CTOPP-2)^29^, which provides an age-adjusted, standardized index of phonological awareness for each participant.

### 2.3 EEG Data Collection During Verb Generation & Resting Conditions

EEG was recorded using a 128-channel saline-based cap (EGI NetStation system) at a 1000Hz sampling rate with electrode impedances maintained below 50 kΩ. Participants were seated comfortably in a sound-proof booth in front of a computer monitor. Children performed a verb generation task, which reliably engages the language network^30,31^. Each trial began with a baseline in which children looked at a central fixation cross; children maintained central fixation throughout the trial to minimize eye movements. There was then an auditory presentation of a noun (500– 1000 millisecond duration), selected from the Children’s Printed Word Database^32^, to which participants were instructed to silently generate an associated verb. Covert response reduces motion artifacts. After three seconds, participants were prompted to vocalize the verb aloud, ensuring task engagement^30,31^. Participants completed two blocks of 60 trials. Following the verb generation task, participants completed a resting-state recording, with alternating blocks of 3-minute eyes open and 3-minute eyes closed.

### 2.4 EEG Data Preprocessing

EEG data was preprocessed using EEGLAB^33^ running in the MATLAB environment and the MADE pipeline^34^, designed for pediatric EEG. Data were down-sampled to 500 Hz, then high-pass (1 Hz) and low-pass (50 Hz) filtered. Bad channels were removed using the FASTER algorithm^35^. Independent Component Analysis (ICA) isolated and removed artifacts, including eye blinks and muscle noise^36^. For verb generation data, EEG was segmented into 3-second epochs beginning at noun onset. For resting-state data, EEG was divided into 3-second non-overlapping epochs. Residual ocular artifacts were identified and epochs rejected if voltage exceeded +/-150uV in electrodes near the eyes. Any epochs with >10% of non-ocular channels exceeding the voltage threshold were also discarded. For remaining epochs, individual channels that surpass the voltage threshold are interpolated at the epoch level. Channels rejected at the beginning of pre-processing were then additionally interpolated. All data were re-referenced to a common average. Finally, a board-certified epileptologist (FMB) visually inspected all epochs to identify and exclude those containing IEDs, as IEDs transiently and dramatically increase functional connectivity^10^ and are not present in controls. This was done to ensure that analyses reflected connectivity related to language processing rather than epileptic activity.

### 2.5 Regions of Interests (ROIs)

Language processing involves a broad range of brain regions including frontal^26,27^, occipitotemporal^28,37^ and motor^38^ cortices. In SeLECTS, seizures and IEDs originate in sensorimotor regions, and these discharges have been associated with both hyperconnectivity and language dysfunction^24^. We aimed to include a broad set of brain regions to capture network dynamics relevant to both language processing and SeLECTS pathology. Accordingly, we defined ten regions of interest (ROIs), five per hemisphere: superior frontal, inferior frontal, motor, temporal, and occipital (Fig. 1A). Each ROI comprised clusters of electrodes corresponding to the underlying cortex, defined according to standard electrode positions^39,40^.

**Figure 1.**
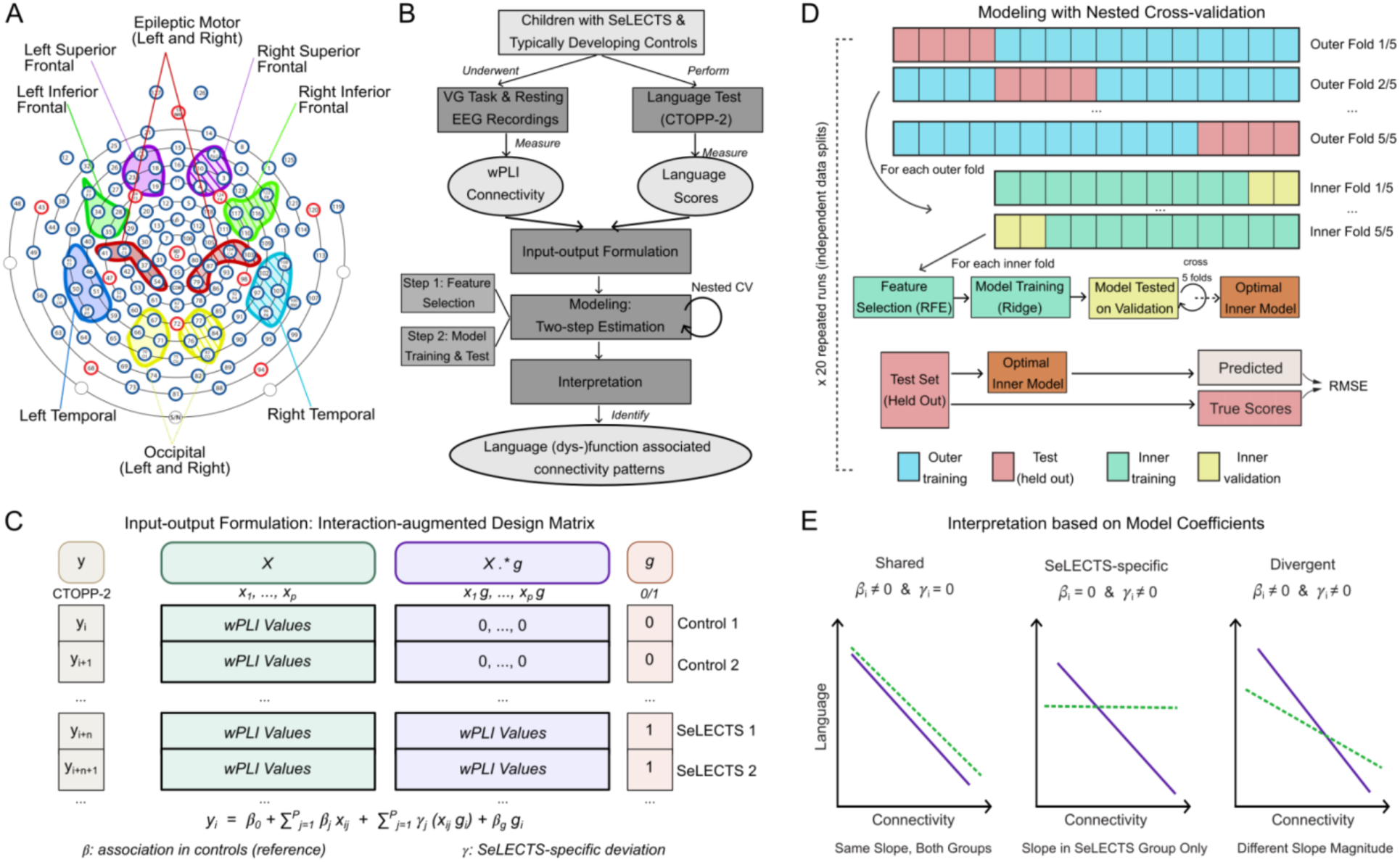
Region of interests (ROIs) & Multivariate pattern analysis (MVPA) pipeline. **A:** ROIs included left and right hemisphere inferior frontal, superior frontal, motor, temporal, and occipital regions. **B**: Flowchart showing the MVPA procedure of identifying language function associated connectivity in children with and without epilepsy. VG Task = Verb Generation Task. **C**: Interaction-augmented design matrix. For each subject *i*, wPLI connectivity values across all unique ROI pairs and four frequency bands were arranged as primary features (X) and group-modulated interaction features (X·g), alongside a binary group indicator (g; SeLECTS = 1, Control = 0) and the outcome vector (y; CTOPP-2 scores). Controls have zeros in the interaction columns by definition. The model equation (below) follows a control-normative formulation in which βj captures the baseline connectivity-language association (estimated in controls as reference group) and γj captures the SeLECTS-specific deviation from that baseline. **D**: Procedure of the nested cross-validation. The dataset is split into 5 folds; in each fold one subset serves as the held-out test set (pink) and the remaining four serve as the training set (blue). Each training set is further divided into 5 inner folds, with each fold containing 4 inner training (green) and 1 validation (yellow) set. Within each inner training set, recursive feature elimination (RFE) selects predictive connectivity features, and a ridge regression model is trained; this model was tested on the validation set. From the 5 models trained across the 5 inner folds, the optimal inner model (orange) was selected and was applied to the held-out test fold to compute prediction error (RMSE). This entire nested CV process was repeatedly run 20 times to obtain an averaged modeling performance. **E**: Connectivity pattern categories interpreted based on model coefficients. Shared patterns (βj≠0, γj=0): the connectivity-language association is present in both groups with the same slope. SeLECTS-specific patterns (βj=0, γj≠0): the association is present in SeLECTS only, with no relationship in controls. Divergent patterns (βj≠0, γj≠0): both groups show a connectivity-language association but in a qualitatively different relationship in SeLECTS relative to controls.

### 2.6 Connectivity Measurements

We used the debiased wPLI to assess connectivity between ROIs. wPLI is a phased-based metric robust against volume conduction, making it suitable for detecting functional connectivity using scalp EEG^25^. Children with SeLECTS have increased wPLI connectivity in several frequency bands compared to controls^18,19^, so wPLI was calculated across four frequency bands (theta [4-7 Hz], alpha [8-12 Hz], beta [13-30 Hz], gamma [31-50 Hz]) between all unique electrode pairs using cross-spectral density within MATLAB-based Fieldtrip’s connectivity toolbox^41^. Including multiple bands allowed us to capture a comprehensive spectrum of connectivity dynamics. To enhance the signal-to-noise ratio, electrode-level wPLI values were averaged to yield **regional connectivity** values representing connectivity between ROIs. For example, theta band wPLI values between all electrode pairs included in the left inferior frontal [LIF] and right superior frontal [RSF] were averaged to produce a regional connectivity (*θ*-LIF-RSF). This procedure resulted in 180 unique regional connectivity values (45 per frequency band).

Children with SeLECTS exhibit widespread increases in functional connectivity compared to typically developing controls^6^, suggesting that hyperconnectivity may reflect a generalized rather than regionally specific phenomenon. To test whether there is regional specificity to connectivity-language associations, we calculated three average connectivity values: **global connectivity (**mean wPLI across all ROI pairs, yielding one per frequency band); **intrahemispheric connectivity (**average across ROI pairs within each hemisphere, yielding 2 values per frequency bands); and **interhemispheric connectivity (**averaged across ROI pairs connecting the left and right hemispheres, yielding one value per frequency band).

### 2.7 Multivariate Pattern Analysis

MVPA was applied to identify language-associated connectivity patterns and to evaluate whether regional connectivity predicts individual language ability beyond averaged connectivity measures or clinical factors. The MVPA pipeline consists of three major stages: input-output formulation, modeling, and interpretation (Fig.1B-E).

#### 2.7.1 Input-output Formulation (Fig.1C)

For each subject *i*, we defined *y_i_* as the language score; *g_i_* as a binary group indicator (SeLECTS = 1, Control = 0), and *X_i_* = [*x_i1_*, . . ., *x_iP_*] as the *1* × *P* feature vector representing all included regional connectivity features (aforementioned 180 wPLI values). To allow connectivity-language associations to differ between groups, we constructed an **interaction-augmented design matrix** consisting of primary connectivity features *x_i_*_j_, group-modulated interaction features *x_i_*_j_ *g_i_*, and the group indicator *g_i_*. This yields the linear model family:

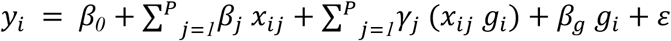

This input-output formulation supports a **control-normative + SeLECTS-deviation** interpretation. As such, for each regional connectivity feature *j*:

- *β*_j_ represents the connectivity-association in controls (reference group).
- *γ*_j_ represents the SeLECTS-specific deviation from the control association. Therefore:
- *β*_j_ + *γ*_j_ represents the connectivity-language association in SeLECTS.

These components yielded a final interaction-augmented design matrix **[*X, X .* g, g*]** with a dimensionality of *N* × (2*P* + 1) as model input, and the corresponding output vector *y* represents language scores (*N* × 1).

This primary MVPA model encodes the hypothesis that connectivity-language associations differ between groups through the inclusion of group-modulated interaction terms. To evaluate whether this assumption is warranted, we fit an otherwise identical MVPA pipeline with primary connectivity features only (without interaction terms) to test whether allowing for SeLECTS-specific associations provides incremental predictive value over a single set of connectivity features applied uniformly to both groups (Supplementary Materials #1).

#### 2.7.2 Modeling (Fig.1B, D)

The modeling stage consists of a two-step estimation. To obtain unbiased estimates of model generalizability, we implemented a five-fold nested cross-validation procedure^42–44^. In each of the five outer folds, one subset of subjects was held out as the test set and remained completely independent during the entire two-step estimation, while the remaining training data was used for feature selection and model training as described below. To minimize variability due to random partitioning, the entire nested cross-validation process was repeated 20 times with different data splits.

##### First step – Feature selection

To identify predictive connectivity patterns while controlling model complexity and thereby reducing the risk of overfitting given the small sample size and large feature space, we applied recursive feature elimination (RFE)^45^ to the full interaction-augmented design matrix. RFE iteratively fits a regularized predictive model, ranks predictors by their contribution to prediction, and removes the least informative predictors until an optimal subset is obtained. We used a generalized linear model with *L*2 regularization (ridge regression^46^) within RFE to stabilize feature ranking under collinearity among connectivity features.

##### Second step – Model training and testing

Features retained by RFE were used to train a second-stage ridge regression model to predict language scores. Within the training data, an inner five-fold cross-validation was performed for feature selection and hyperparameter tuning. Predictive accuracy was quantified using the root mean square error (RMSE) of language score predictions on the held-out test set. This separation ensures that the reported performance reflects true out-of-sample prediction^42,43^. To facilitate efficient implementation of the MVPA pipeline, parallel computing strategies were employed, assigning each run as an independent task distributed across separate computing cores^47^.

This two-step estimation framework enabled us to identify and validate the connectivity patterns most strongly associated with language function.

#### 2.7.3 Interpretation (Fig.1E)

Three complementary metrics were used to interpret the model. First, **model coefficients** classified each retained feature into one of three categories based on which terms RFE selected: *Shared patterns* (nonzero *β*_j_, zero *γ*_j_), indicating a baseline connectivity-language association present across both groups; *SeLECTS-specific patterns* (zero *β*_j_, nonzero *γ*_j_), indicating an association present only in SeLECTS; and *Divergent patterns* (nonzero *β*_j_ and *γ*_j_), indicating that the association differs in magnitude or direction between groups. Coefficients reflect the direction and strength of connectivity-language associations within the model but do not directly quantify predictive contribution, particularly under regularization and in the presence of correlated features. **Permutation feature importance (PFI),** our second metric, addresses this by estimating each feature’s out-of-sample predictive contribution: feature values were permuted 1000 times and the resulting mean change in RMSE taken as the PFI. Features with larger PFI have greater influence on out-of-sample prediction, providing information complementary to model coefficients. Third, **weighted single-region importance** translates connection-level PFI into a region-level summary. Each connection’s PFI was distributed equally to its two constituent regions, summed across all retained connections involving that region, and normalized to percentages. We performed this step because wPLI is an undirected measure and does not attribute importance to one region over the other.

### 2.8 Analytic Strategy and Statistical Comparisons

Four analyses evaluated our hypotheses about connectivity-language associations, all using the MVPA procedure described above. Verb generation and resting EEG were modeled separately, as connectivity during both task and rest has been associated with language in SeLECTS^9,10^. The first analysis tested whether regional connectivity predicts language above chance, comparing model performance against a null model that predicted the mean language score across all participants. The remaining three analyses tested whether regional connectivity provides incremental predictive power, each comparing a model containing a simpler set of predictors (averaged connectivity or clinical variables) against a model additionally incorporating regional connectivity. Model performance was quantified as RMSE and compared using Wilcoxon signed-rank tests on distributions across the 20 repeated runs. Table 2 summarizes model structure and performance.

#### 2.8.1 Main Model: Regional Connectivity

We fit the MVPA using the full interaction-augmented design matrix to predict language scores, separately for task and resting EEG, comparing each to the null model. We also compared task- and resting-based models to each other to assess the relative informativeness of EEG condition (3 comparisons, Bonferroni corrected threshold p<0.017). We visualized the anatomical distribution of Shared, SeLECTS-specific, and Divergent connectivity patterns by ranked PFI. To provide directional intuition about top-ranked connections, we report univariate Spearman’s correlations between individual regional connectivity values and language scores, but these are descriptive and not subjected to formal hypothesis testing.

#### 2.8.2 Regional vs. Averaged Connectivity

To test whether regional connectivity offers predictive advantage over simpler summary measures, we compared a model fit using all three averaged connectivity measures combined (global, intrahemispheric, and interhemispheric) against a model additionally incorporating regional connectivity, separately for task and resting EEG (2 comparisons, corrected threshold p<0.025). Models fit with single averaged connectivity measures are visualized in Supplementary Figure #3. As a model with 180 regional features would be expected to outperform one with 3 summary features on purely statistical grounds, we also examined whether any averaged connectivity feature was retained among the top-ranked predictors in the regional model.

#### 2.8.3 Connectivity vs. Clinical Factors

To test whether regional connectivity predicts language ability beyond clinical information, we compared models fit with clinical variables alone (epilepsy diagnosis, ASM use, age, and all three combined) against models additionally incorporating regional connectivity, separately for task and resting EEG (8 comparisons; corrected threshold p<0.00625).

#### 2.8.4 Clinical Validity: Prediction in Lower-Scoring Children

The primary MVPA model optimizes prediction accuracy uniformly across the full language score distribution, making it well-suited to characterize connectivity-language associations across the population. However, lower-scoring children are the most clinically relevant target for connectivity-guided neurostimulation. Running a weighted model as the primary analysis would risk optimizing for this subgroup at the expense of broader predictive validity and generalizability. We therefore used a weighted variant of the MVPA as a validation analysis in which we deliberately amplified the influence of lower-scoring children during training (by assigning them a higher *observation weight*) with the goal of testing whether the connectivity patterns identified by the primary model (learned without that bias) remain predictive in this clinically prioritized subgroup. Both models used identical input features and differed only in whether observation weights were applied. Full methodology details of this weighted version are available in Supplementary Material #2.

We compared the weighted and primary MVPA on three outcomes. *Prediction accuracy* was assessed using both standard RMSE (equal weighting across participants) and weighted RMSE (errors scaled by observation weights), enabling comparison of overall and lower-score-specific accuracy. *Classification of lower-scoring children* was assessed using a quadrant-based analysis. Lower-scoring children were defined as those scoring more than 1 SD below the observed sample mean on the CTOPP-2 scores, a threshold that exceeds the test’s reported test-retest variability of approximately 2–3 points^48^ and identifies genuine underperformance while remaining feasible given SeLECTS’ association with mild-to-moderate rather than severe phonological impairment^14,23^. As such, subjects were classified as correctly identified lower-scoring (both observed and predicted scores below threshold), missed lower-scoring (observed below, predicted above threshold), or non-lower-scoring. *Convergence of selected features* was assessed qualitatively by comparing the connectivity features retained as top predictors across the primary and weighted MVPA. Strong overlap indicates the primary model identified patterns reflecting connectivity genuinely relevant to lower-scoring children.

## 3. Results

### 3.1 Participants, Language Scores & Data Quality

Thirty-six children with SeLECTS and 32 age-matched typically developing controls were consented. Nine children with SeLECTS and three controls were excluded for either poor EEG quality (three SeLECTS, one control) or failure to complete the CTOPP-2 (six SeLECTS, two controls). There were no significant group differences in age or sex in the final cohort (Table 1). Sixteen (59%) SeLECTS participants were taking ASM: 9 levetiracetam, 7 oxcarbazepine. Significant differences in CTOPP-2 scores were observed between the two groups (SeLECTS 102+/-14; controls 113+/-14; *p* = 0.001). There were no significant differences in data quality and retained trials after preprocessing (Supplementary Table #1).

**Table 1:** Demographics & cognitive scores for children with SeLECTS and controls. P-values are from two-sample t tests compared between groups (SeLECTS vs. Controls).

| Total (n=56) | SeLECTS (n=27) | Controls (n=29) | p-value |
| --- | --- | --- | --- |
| Age | 10.1±1.7 | 9.8±1.8 | 0.52 |
| Sex (male), n (%) | 19(70) | 16 (55) | 0.24 |
| Medication, n (%) | 16 (59) | N/A | N/A |
| CTOPP-2 scores | 102.1±13.8 | 113.2±14.1 | 0.004 |

### 3.2 Predicting Language from Regional Connectivity

Comparing to the null model (RMSE=14.78), both the task (RMSE=10.88±0.62; *p*<0.001) and resting-state (RMSE=12.53±1.03; *p*<0.001) models yielded significantly better predictions, with the task model also outperforming resting-state (paired p<0.001). Supplementary models using only primary connectivity features without interaction terms performed significantly worse than models with interaction features (task RMSE=12.411±0.646; resting RMSE=14.174±0.930), demonstrating that the SeLECTS group has connectivity-language patterns unique from those of controls (Supplementary Materials #1).

### 3.3 Group-shared and SeLECTS-specific Patterns

Recursive feature elimination identified 36 connectivity features for task EEG and 32 features for resting EEG. Feature selection was stable across all nested CV folds and repeated runs, demonstrating that the retained features represent robust, partition-independent predictors of language ability rather than artifacts of a particular data split.

#### Connectivity During Verb Generation Task

Of the 36 retained task EEG features, 15 were classified as Shared, 15 as SeLECTS-specific, and 6 as Divergent patterns. We ranked these features according to the mean of their PFI values and examined their anatomical distribution. Shared patterns predominantly involved frontal, temporal, and occipital connections in the left hemisphere, whereas motor-related connections were uniquely predictive in SeLECTS, consistent with the epileptic onset zone (Fig.2B and Fig.3). Consistent with this pattern, weighted single-region importance analysis showed that the shared feature set was dominated by left-hemisphere regions: the left superior frontal region accounted for 23% of the total importance, followed by the left occipital (18%), left temporal (15%), and right inferior frontal (14%) regions. In contrast, for SeLECTS-specific features, the left motor region contributed the largest proportion of the weighted importance (20%), with additional contributions from the right occipital (19%), right temporal (12%), and right motor (11%) regions.

**Figure 2:**
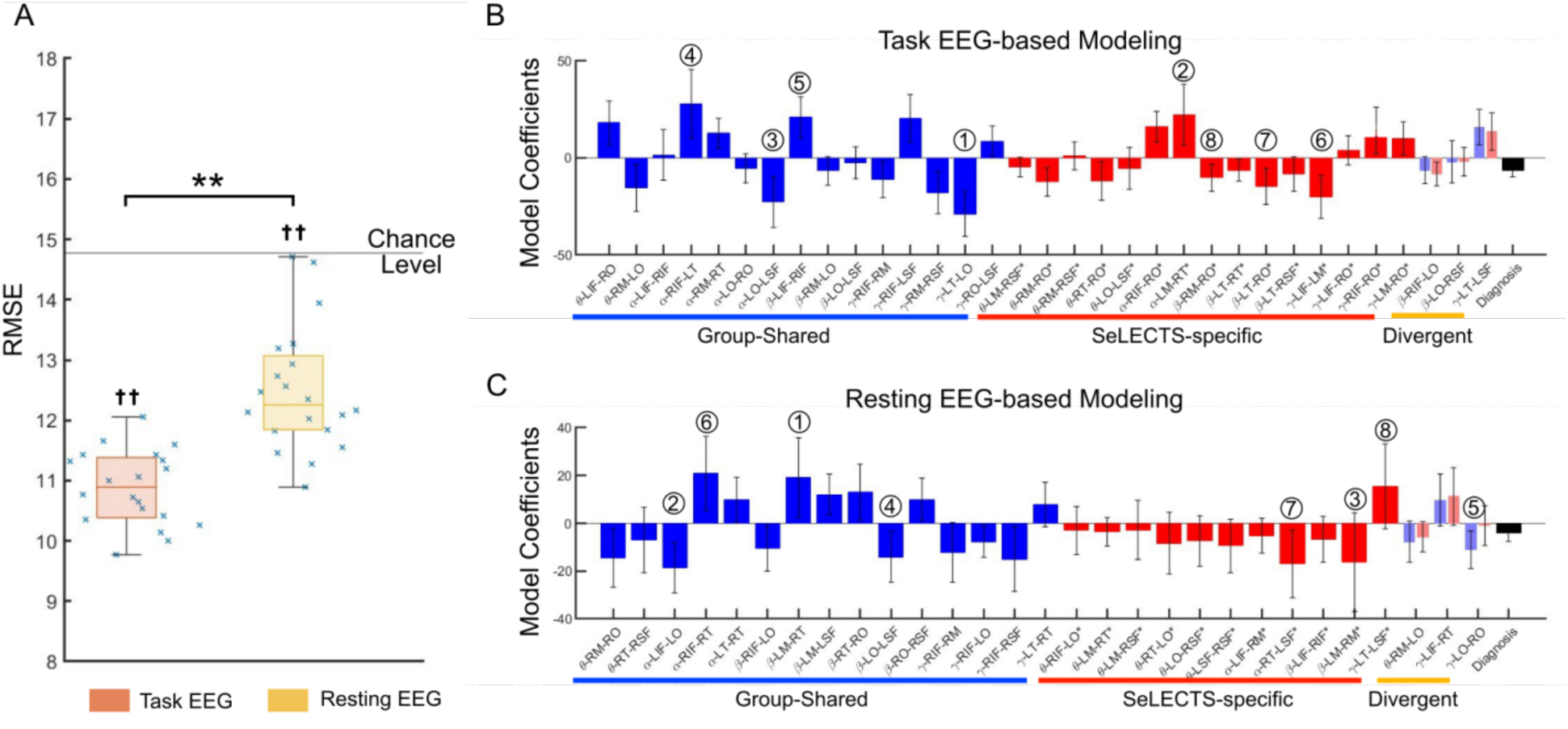
**A**. Multivariate pattern analysis (MVPA) predicted language scores from both task-based and resting-state EEG connectivity significantly better than chance, quantified by root mean squared error (RMSE). Each cross represents the RMSE from a single run. Orange boxplots show the RMSE distribution for models using task EEG, and yellow boxplots show the RMSE distribution for models using resting EEG. †† represents significant after multiple comparison comparing to the chance level; and ∗∗ represents significant after multiple comparison correction in the paired comparisons. B and C. Model coefficients for connectivity patterns identified by MVPA using task (B) and resting EEG (C). Blue bars indicate shared patterns, and red bars indicate SeLECTS-specific patterns. For divergent patterns, light blue indicates control patterns (*β*_j_) and light red indicates SeLECTS-patterns (*γ*_j_). Black bar indicates the group indicator. Error bars represent the standard deviation of coefficients across the 20 repeated runs. The eight most important features, sorted by their PFI values are annotated for illustration purposes.

**Figure 3:**
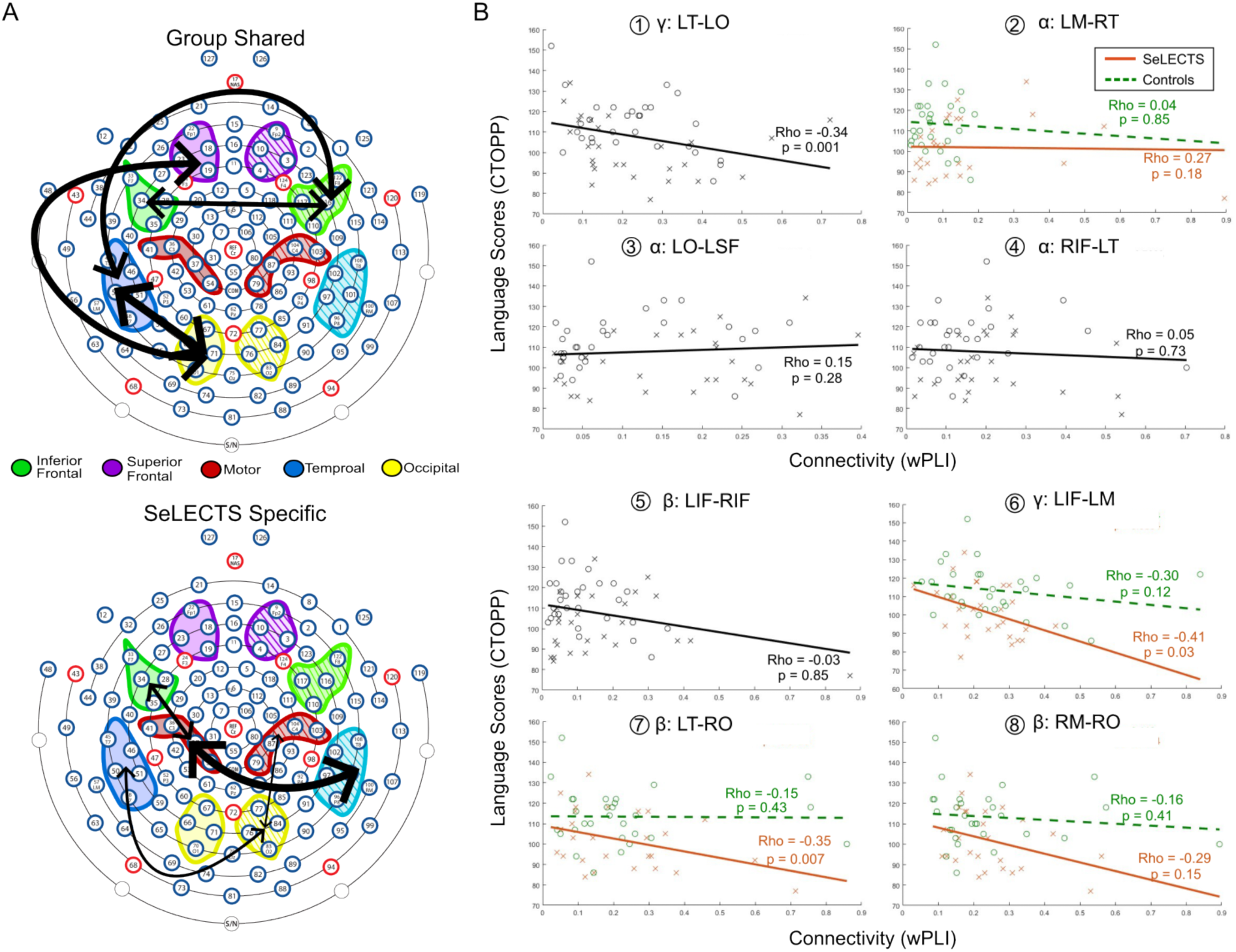
Language-associated connectivity patterns from <u>task EEG</u>. **A**: Model-identified important language-associated connectivity patterns which are <u>shared</u> (top) and <u>SeLECTS-specific</u> (bottom). Black arrows represent connectivity between two regions, with arrow weight indicating the quantified importance of that connection in predicting language. **B**: Correlation between language function and regional connectivity in model-identified important features visualized in (A). P-values are derived from Spearman’s correlation, while the lines are fitted using a linear regression model for illustrative purposes (one line for shared patterns; individual lines for SeLECTS-specific patterns). Sub-plots are ordered based on their PFI ranking. Crosses represent values from SeLECTS while circles represent values from controls. Region labels follow the format L or R for hemisphere and IF = inferior frontal, SF = superior frontal, M = motor, T = temporal, O = occipital.

All three interpretation variables (model coefficients, PFI values and weighted single-region importances) were highly consistent across models, irrespective of the inclusion of clinical covariates (Supplementary Figure 4-5), indicating that the MVPA yields stable and meaningful feature selection.

Univariate Spearman correlations at the individual connection level showed a consistent trend across the top-ranked retained connections. Higher connectivity was associated with lower language scores, particularly in SeLECTS (Fig.3B). Most individual connections did not reach corrected significance, consistent with the distributed, multivariate nature of the connectivity-language signal and with the rationale for the MVPA approach.

#### Connectivity During Rest

Of the 32 retained resting EEG features, 15 were classified as Shared, 11 as SeLECTS-specific, and 6 as Divergent patterns. Left frontal, temporal and occipital connections were again identified as shared patterns, whereas motor-related connections emerged as both shared and specific: left motor–right temporal connectivity was shared, whereas bilateral motor connectivity was uniquely predictive in SeLECTS (Fig.2C and Fig.4).

**Figure 4:**
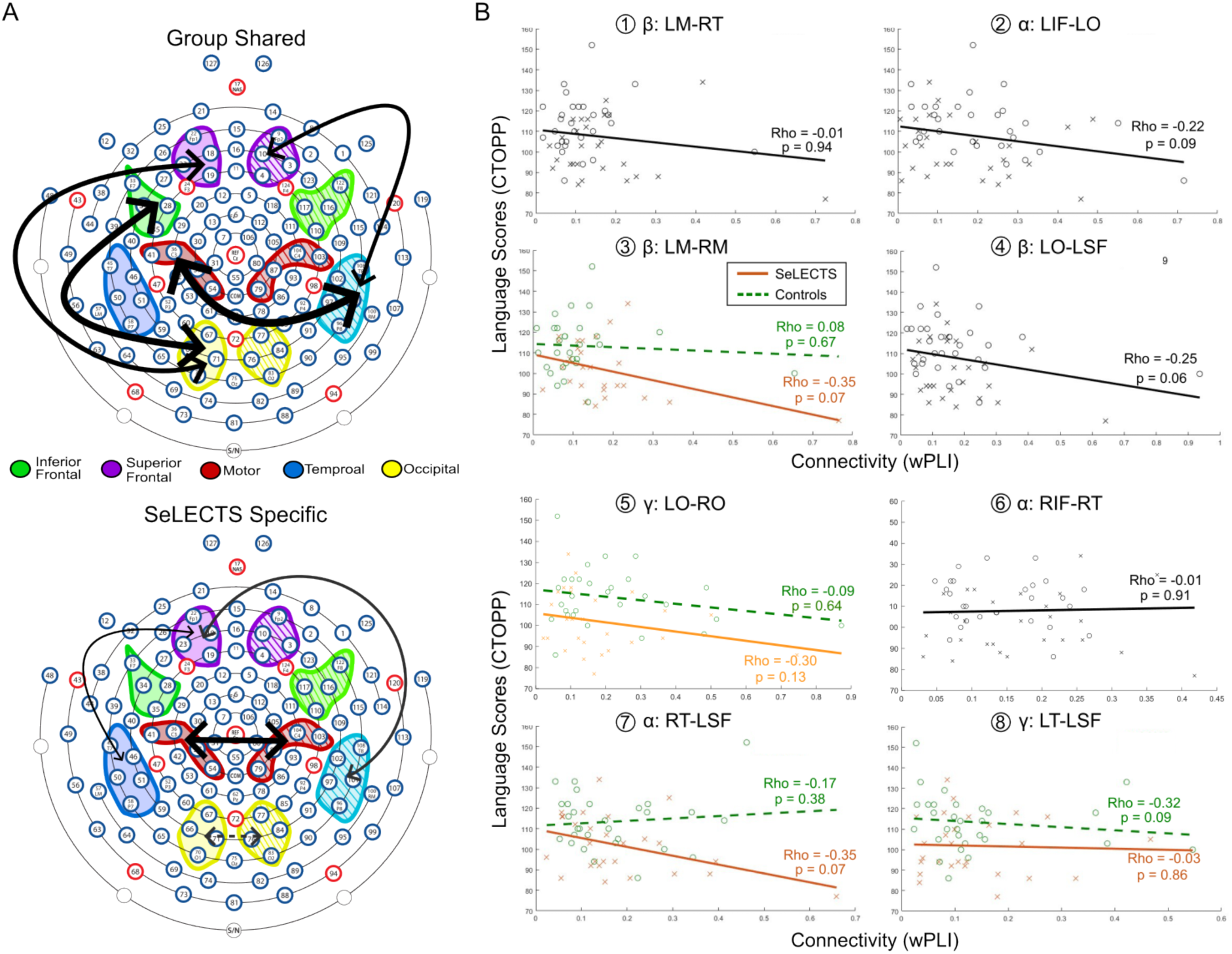
Language-associated connectivity patterns from <u>resting EEG</u>. **A**: Model-identified important language-associated connectivity patterns which are <u>shared (top)</u>, <u>divergent (top, dashed)</u>, and <u>SeLECTS Specific</u> (bottom). Black arrows represent connectivity between two regions, with arrow weight indicating the quantified importance of that connection in predicting language. Notably, the LO-RO connectivity was ranked as divergent, but its SeLECTS-specific feature was with a small importance. **B**: Correlation between language function and regional connectivity in model-identified important features visualized in A. P-values are derived from Spearman’s correlation, while the lines are fitted using a linear regression model for illustrative purposes (one line for shared patterns; two lines for SeLECTS-specific or divergent patterns). Sub-plots are ordered based on their PFI ranking. Crosses represent values from SeLECTS while circles represent values from controls. Region labels follow the format L or R for hemisphere and IF = inferior frontal, SF = superior frontal, M = motor, T = temporal, O = occipital.

For the shared patterns, weighted single-region importance analysis indicated that the left occipital region showed the highest weighted importance (19%), followed by the right temporal (19%), left inferior frontal (12%), and right occipital (11%) regions. For the SeLECTS-specific features, the left superior frontal region contributed most strongly (20%), with additional contributions from the right motor (15%), left motor (15%), and right temporal (11%) regions.

As in the task condition, the quantitative importance of these connectivity features remained stable across models regardless of clinical covariates (Supplementary Figure 6-7).

### 3.4 Regional vs. Averaged Connectivity

Task and resting models incorporating regional connectivity outperformed models relying on average connectivity (Fig.5, Table 2). All average connectivity features performed similarly to each other and none of them were retained by the RFE step, meaning not enough predictive power to language compared to the regional connectivity.

**Figure 5:**
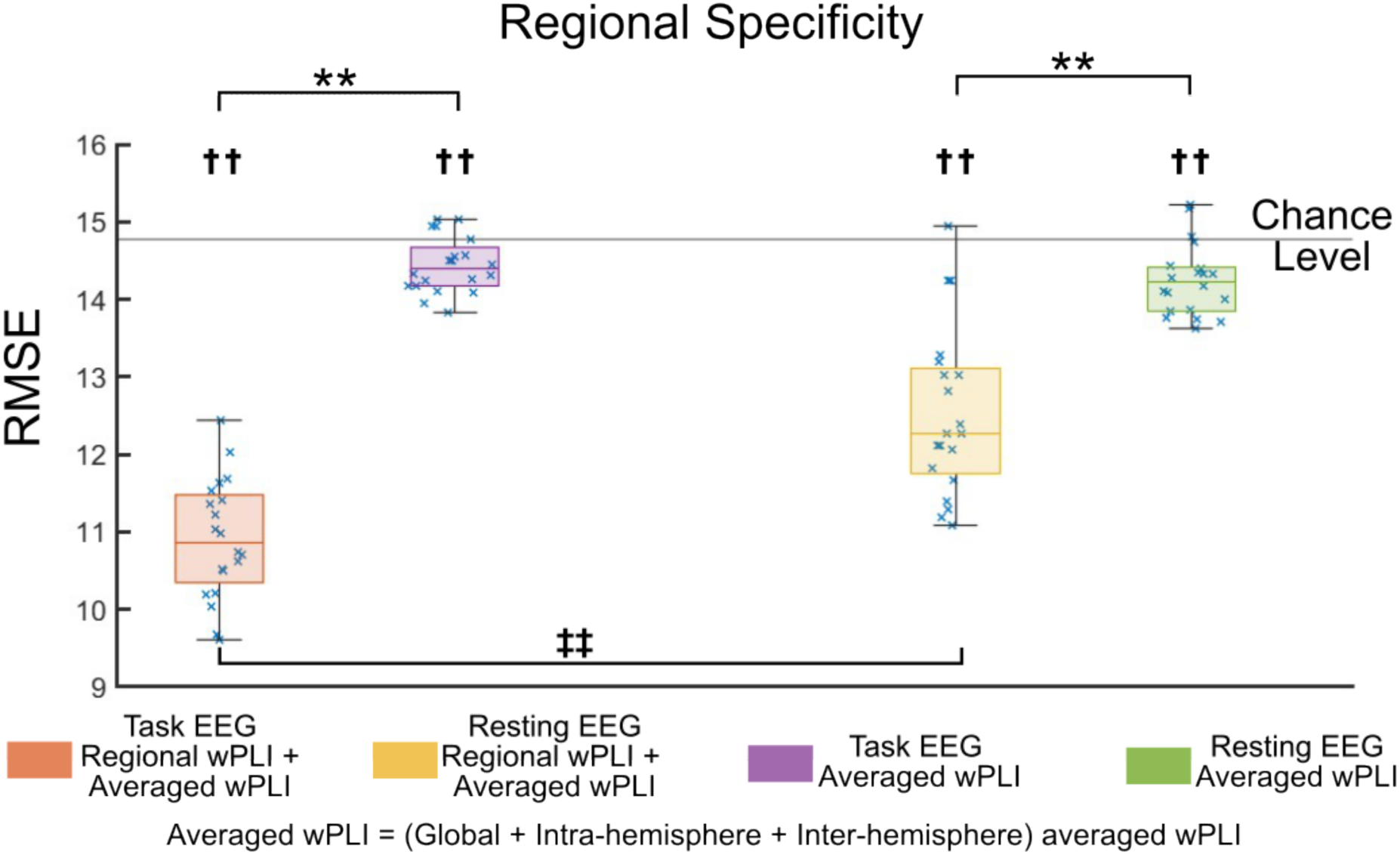
Prediction performance, measured by root mean squared error (RMSE), of the tested models. Each cross represents the RMSE from a single run. Box plots depict the distribution of these performances. Orange boxes: model inputs consist of regional connectivity calculated from <u>task</u> EEG combined with the averaged measures; purple boxes: model input was the averaged connectivity measures from <u>task</u> EEG alone; yellow boxes: model inputs consist of regional connectivity calculated from <u>resting</u> EEG combined with the averaged measures; green boxes: model input was the averaged connectivity measures from <u>resting</u> EEG alone. †† represents significant after multiple comparison correction comparing to the chance level; and ∗∗ represents significant after multiple comparison correction in the paired comparisons. ‡‡ represents significant after multiple comparison correction in the paired comparison between task and resting EEG based connectivity.

**Table 2.**
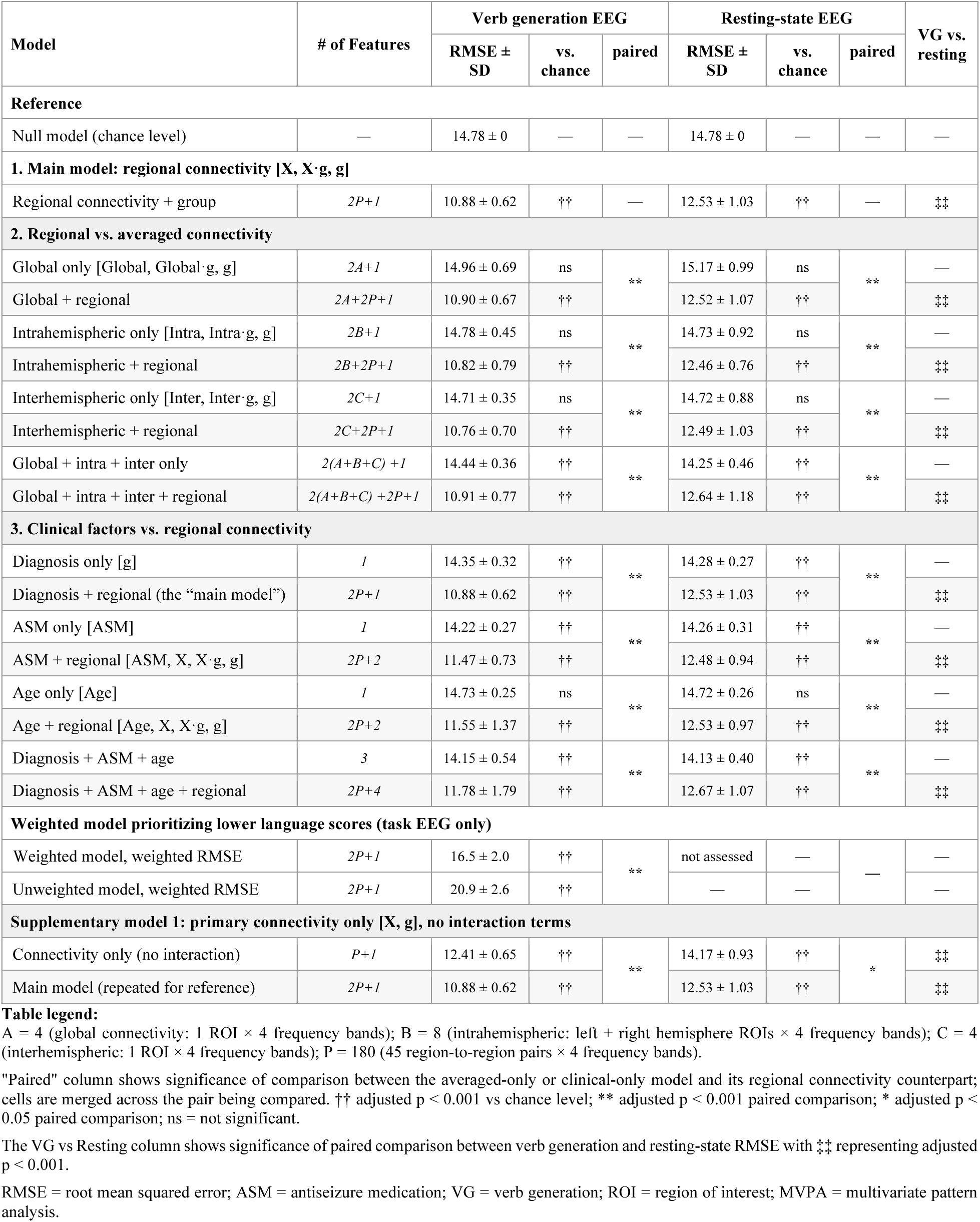
Model performance summary: prediction of language scores (RMSE) across all tested models.

### 3.5 Clinical Factors

Age did not outperform chance, consistent with the fact that CTOPP-2 is age-normalized. Models incorporating epilepsy diagnosis or ASM use outperformed chance and were themselves significantly outperformed by models including task or resting regional connectivity (Fig.6, Table 2).

**Figure 6:**
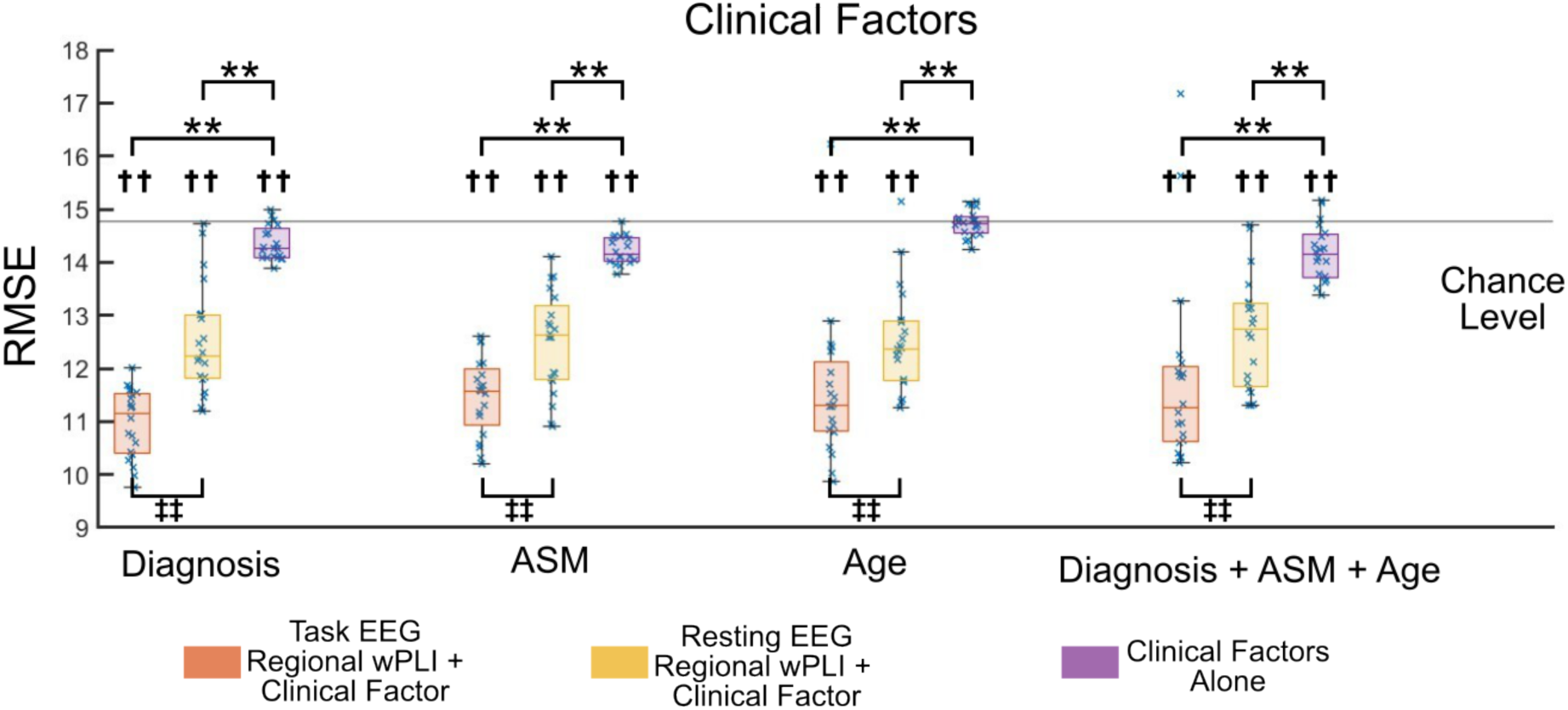
Prediction performance, measured by root mean squared error (RMSE), of the tested models. Each cross represents the RMSE from a single run. Box plots depict the distribution of these performances. Orange boxes: model inputs consist of regional connectivity calculated from <u>task</u> EEG combined with investigated clinical factors (i.e., diagnosis, ASM use, age); Yellow boxes: model inputs consist of regional connectivity calculated from <u>resting</u> EEG combined with the investigated clinical factors; purple boxes: model input was the clinical factors alone. †† represents significant after multiple comparison correction comparing to the chance level; and ∗∗ represents significant after multiple comparison correction in the paired comparisons. ‡‡ represents significant after multiple comparison correction in the paired comparison between task and resting EEG based connectivity.

### 3.6 Clinical Validity: Prediction in Lower-Scoring Children

The weighted and primary MVPA models were compared on two accuracy metrics to confirm that upweighting lower-scoring children during training meaningfully shifted model behavior. Standard RMSE assessed overall prediction accuracy across the full cohort; weighted RMSE assessed prediction accuracy across all participants but assigned greater weight to errors in lower-scoring children. The weighted MVPA improved weighted RMSE relative to the primary model (16.5±2.0 vs. 20.9±2.6, p<0.001), confirming that the weighting procedure successfully prioritized lower-scoring children, at a small cost to overall accuracy as measured by standard RMSE (weighted MVPA 11.7±0.9 vs. primary MVPA 10.9±0.6, p=0.02). In the quadrant-based classification, the weighted MVPA correctly identified 5.8±1.4 out of 9 lower-scoring children compared with 3.9±0.1 for the primary model (p<0.01) (Figure 7B-C). Even with this shift in model behavior, the two models identified a largely overlapping set of connectivity patterns, including both shared and SeLECTS-specific predictors (Fig.7D-E), indicating that the primary MVPA identified connectivity-language associations robust in lower-scoring children.

**Figure 7:**
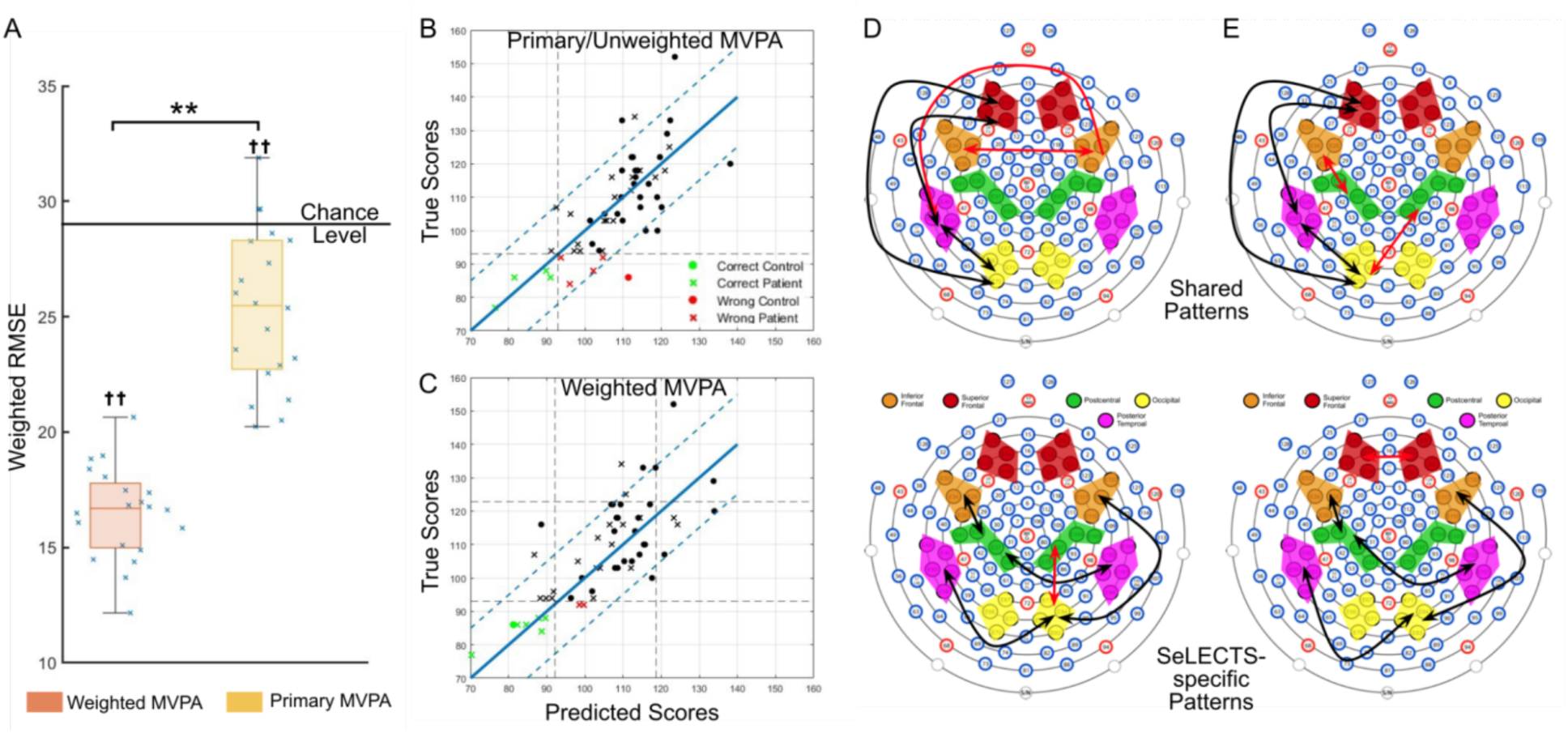
**A**: Prediction performance from the weighted MVPA (orange box, hyperparameter *strength* = 1) versus the primary MVPA (yellow box, *strength* = 0 [unweighted]). Each cross represents the RMSE from a single run. **B and C**: Representative runs (run with lowest inner validation RMSE) of quadrant-based analysis comparing the unweighted/primary vs. weighted MVPA method, illustrating the weighted MVPA correctly identified more lower-scoring children. Green points: corrected classifications; red points: wrong classifications; crosses: children with SeLECTS; filed circles: controls. **D**: Shared (top) and SeLECTS-specific (bottom) connectivity patterns identified by the primary MVPA. **E**: Shared (top) and SeLECTS-specific (bottom) connectivity patterns identified by the weighted MVPA. Black arrows indicate the overlapped connections between the weighted and primary MVPA methods while red arrows indicate connections unique to one model.

## 4. Discussion

This study establishes functional connectivity derived from high-density EEG as a powerful, quantitative predictor of language ability in children with and without epilepsy. Using MVPA, we demonstrated that regional connectivity patterns predict individual language ability at the single-subject level, adding significant predictive power beyond conventional clinical variables including epilepsy diagnosis and ASM use known to be associated with language outcomes. Regional specificity and task engagement proved by spatially-resolved regional connectivity patterns outperformed global averages, and connectivity measured during active language engagement surpassed resting-state measures, revealing that language-relevant brain dynamics are most apparent under cognitive demand. Connectivity patterns shared by controls and children with SeLECTS involve largely canonical left-hemisphere language regions while SeLECTS-specific connectivity patterns centered on motor cortex connectivity, implicating the epileptogenic zone in language dysfunction. These findings establish regional, task-based connectivity as a mechanistically-informative biomarker that captures variation in language function beyond what can be inferred from clinical factors alone.

In both groups, higher connectivity was generally associated with worse language. This pattern would not have been readily apparent from traditional univariate analyses because many individual connectivity pairs showed non-significant correlations with language scores, likely because language processing relies on distributed, interactive networks rather than isolated regional connections^49^. The MVPA approach instead evaluates combinations of connectivity features simultaneously, capturing multivariate patterns where the specific configuration and interaction of multiple regional connections collectively predict language scores, even when isolated individual connections show weak effects^20,50,51^. The association between high connectivity and poor language may reflect failed network segregation. In typical development, language networks become increasingly left-lateralized and functionally segregated from homologous right hemisphere regions^52^. Children with epilepsy who display reduced intra-network integration and excessive, diffuse internetwork coupling have worse cognitive outcomes^53^. IEDs may drive some of these network issues, leading to both acute^6,7,54^ and more persistent connectivity between epileptic and language cortices^18,19^. While higher connectivity could represent compensatory network plasticity, wherein regions outside of the typical language nodes are recruited to support language function^11^, this is less likely as we find that the relationship between connectivity and language is almost always negative.

### Shared Connectivity Patterns

Shared patterns largely favored left hemisphere connections, consistent with the typical left-lateralized language network^26–28^. Prior fMRI studies show divergent findings regarding the role of left hemisphere connectivity in children. A task-based study during story listening demonstrated similar connectivity in language networks compared to controls^55^, whereas a resting-state fMRI study found reduced left-hemisphere connectivity and loss of the connectivity-language relationship seen in controls^56^. We find converging patterns in both task and resting data, where higher left hemisphere connectivity is associated with worse language in both groups. Our multivariate approach, which simultaneously evaluates connectivity patterns across multiple regions, may have revealed these relationships that were not apparent in prior univariate analyses, suggesting that high connectivity (at least as measured by EEG) reflects inefficient network organization.

### SeLECTS-specific Connectivity Patterns

Motor region connectivity emerged as a critical feature associated with language outcomes in the SeLECTS group, especially during the language task. IEDs in SeLECTS acutely enhance functional connectivity between motor and language regions, with connectivity strength negatively correlating with performance IQ^7^. Motor cortex involvement may cause dysfunction at both local and network levels. At the local level, the primary motor cortex has recently been found to directly participate in phonological discrimination through auditory-like representations^57,58^, such that epileptic activity may directly disrupt phonological processing. At the network level, aberrant motor-language coupling may prevent the functional segregation characteristic of typical development^53^, interfering with efficient language processing. These complementary mechanisms may together account for both the phonological deficits we observed and the broader language impairments characteristic of SeLECTS.

### Regional vs. Averaged Connectivity

Regional connectivity patterns consistently outperformed global and hemispheric averages, demonstrating that the language-relevant signal in this population requires spatial resolution to be captured. This finding argues against the interpretation that diffuse hyperconnectivity alone explains language impairment in SeLECTS. If global elevation were the primary driver, averaged measures would perform comparably to regional ones. Prior work supports this regional specificity. In pediatric focal epilepsy, high temporal connectivity was associated with worse language while high frontal connectivity was associated with better language^9,10^, such that averaging would obscure both. The translational implication is that spatially targeted neurostimulation approaches may outperform broader ones when treating cognitive comorbidities. Data from responsive neurostimulation (RNS), which targets identified seizure foci, support this. RNS shows a favorable cognitive profile compared to ablative approaches^59^, and benefits depend on the region implanted; improvements in verbal learning occurred in patients with mesial temporal leads, while naming gains were observed in those with neocortical leads^60^. Furthermore, pre-implant regional, but not global, functional connectivity predicts RNS response, supporting the principle that spatially resolved connectivity measures should guide stimulation site selection for future interventional studies^61^.

### Clinical Features

A critical question is whether connectivity captures meaningful and unique individual variation in language, or whether it simply reflects what a clinician could surmise based on clinical factors alone. Children with SeLECTS on average perform worse on language tests than typically developing children, with medicated children with SeLECTS performing worse than unmedicated peers^62^. Furthermore, both SeLECTS itself and ASM use have been associated with alterations in functional connectivity^3,14^. As such, we wanted to ensure that connectivity was not simply a proxy for diagnosis or medication use. In our models, age added no predictive power, consistent with age-normalization of CTOPP-2, while diagnosis and ASM use (individually or together) barely outperformed the null model. In contrast, connectivity added substantial predictive power beyond these clinical variables, suggesting it captures subject-specific network organization beyond clinical history. Children with SeLECTS have significant variability in language outcomes. Here, we find that connectivity distinguishes between them in a way that clinical variables cannot.

### Task vs. Resting-Based Connectivity

Task-based models outperformed resting-based models, though both predicted language meaningfully. This aligns with adult studies showing task-evoked brain states provide greater sensitivity to cognitive processes than resting baselines.^63^ Task connectivity explains over 20% of fluid intelligence variance (vs. <6% from rest)^64^, better predicts activation maps of brain networks^65^, and more reliably predicts working memory in EEG^66^. Our findings extend this principle to pediatric epilepsy and language, suggesting connectivity during active language engagement reveals network dynamics more effectively than resting-state data. Task-based connectivity may therefore be preferable for future clinical applications, though the predictive value of resting-state connectivity suggests it could serve as a useful alternative when task-based recording is not feasible.

### Clinical Implications

Regional connectivity represents a potentially modifiable therapeutic target. In our previous work, low-frequency rTMS effectively modulated both IEDs and hyperconnectivity in children with SeLECTS^12^, demonstrating that the network abnormalities identified here are amenable to noninvasive intervention. Motor-region connectivity emerged as the dominant SeLECTS-specific feature in both task and resting conditions, consistent with the origin of IEDs in sensorimotor cortices and with prior evidence linking motor-language network disruption to language dysfunction in SeLECTS^3,67^. This implicates the motor network as a mechanistically grounded neurostimulation target, and more broadly suggests that the epileptic onset zone may represent a candidate target in other focal epilepsy populations. These connectivity patterns are equally relevant to the children with lowest language scores who are most likely to benefit from intervention. The weighted model, which better predicted language in these lower-scoring children, identified highly overlapping connectivity patterns as the primary, unweighted model, suggesting these connectivity patterns are robust and important in the most affected children. Higher connectivity was consistently associated with worse language, suggesting the observed hyperconnectivity is pathological rather than a compensatory reorganization, though whether reducing connectivity improves language outcomes requires prospective interventional study. Connectivity may also serve as a prognostic biomarker for developmental outcomes in children with epilepsy. Precedent exists, as connectivity models derived from both intracranial^68^ and scalp EEG^69^ successfully predicted epilepsy surgery outcomes. Translating this approach to language outcomes would require longitudinal designs tracking language trajectories in epilepsy syndromes in which children have a high risk for language deterioration over time. Once causal relationships are established, connectivity could identify at-risk children, complement presurgical language mapping, and monitor treatment responses.

### Limitations

First, connectivity was estimated at sensor level, as individualized MRI scans for subject-specific source reconstruction were not available for this pediatric cohort. Although source-space analyses would provide greater anatomical precision, wPLI is highly robust to volume conduction in sensor space^25^, and the consistent outperformance of regional over global averaged connectivity measures supports that meaningful spatial information is preserved at this level of analysis. Second, the small sample size introduced several related constraints. Despite rigorous nested cross-validation with multiple random partitions, regularization, and consistently better-than-chance performance across control analyses with stable feature importance profiles, the risk of overfitting cannot be fully excluded and replication in larger independent cohorts is essential. Sample size also precluded more flexible modeling approaches. Ridge regression was necessary given the high-dimensional, small-sample setting, but linear models cannot capture threshold effects, interactions between regions, or nonlinear relationships between connectivity magnitude and language score that may be biologically meaningful. Larger samples would enable nonlinear and interaction-based approaches that better reflect the complexity of language network organization. Third, by modeling each frequency band as an independent set of connectivity features, the current approach does not capture cross-frequency coupling. Theta-gamma coupling during phonological encoding is a particularly relevant example of a mechanism that operates across rather than within frequency bands^70^. Treating bands as separate predictors may therefore miss components of the connectivity-language signal that arise from coordinated interactions across frequencies. Future work incorporating cross-frequency coupling metrics would address this directly. Fourth, unmeasured variables including socioeconomic status, general cognitive ability, and ASM-specific effects such as drug type, dose, and duration could confound the observed connectivity-language associations if they independently drive both connectivity and language outcomes. Sample size constraints prevented their inclusion or required collapsing them into binary terms. However, a non-neurological confounder would be expected to produce diffuse, anatomically nonspecific effects. The selective emergence of motor connectivity as a SeLECTS-specific predictor, consistent with the epileptic onset zone, is more consistent with disease-related network disruption than with broad demographic or clinical confounding.

## 5. Conclusion

This study shows that spatially specific patterns of functional connectivity predict individual language ability in children with and without epilepsy and provide information beyond conventional clinical variables. More broadly, this study illustrates how a focal neurological disorder can selectively alter components of a distributed cognitive network while leaving other aspects of network organization shared with typical development. Connectivity involving the motor network, the epileptogenic system in SeLECTS, emerged as particularly relevant to language dysfunction, suggesting a mechanistically promising target for future neuromodulation. Longitudinal and interventional studies will be critical for determining whether modifying these disease-related network patterns can causally improve language outcomes.

## Data Availability

The datasets generated and/or analyzed during the current study are not publicly available due to privacy concerns. The data is clinical data that has not been de-identified. But the code with example data used for illustrating the methodologies of connectivity and modeling is available at https://github.com/Pediatric-Neurostimulation-Laboratory/languageConnectivityBiomarker. All source data for reproducing result figures are also available. The full data is available from the corresponding author on reasonable request.

## Acknowledgements

We would like to thank Dr. Emily Fox, Dr. Scott Linderman, Dr. Heidi Feldman, Hyun Dong Lee for their valuable methodological guidance and thoughtful discussions. We also thank Dr. Torsten Baldeweg for sharing the list of words used for the verb generation task in this study.

## Funding Information

XS receives support from the Stanford Maternal & Child Health Research Institute (MCHRI Postdoctoral Support Fellowship). FMB’s work is supported by a K23 Career Development Award (NINDS K23NS116110), the Doris Duke & Rita Allen Foundation, the Wu Tsai Neuroscience Institute, and a gift from the Principe & O’Farrell family.

## Author Contributions

X.S.: conceptualization, methodology, software, formal analysis, data curation, writing– original draft & editing, visualization. O.P.: validation, writing-review & editing. W.Q.: data curation, validation, writing–review & editing. M.M.: data curation, validation, writing-review & editing. K.C.N.: data curation, validation, writing–review & editing. W.W.: methodology, writing–review & editing. Z.H.: methodology, writing–review & editing. F.M.B.: conceptualization, methodology, formal analysis, data curation, validation, writing–drafting, review & editing, project administration, funding acquisition.

## Potential Conflicts of Interests

None of the authors has any conflict of interest to disclose.

## Notes

### Competing Interest Statement

The authors have declared no competing interest.

