## Supplementary Materials for "Language Dysfunction Associated with Pathological Brain Connectivity in Children with Epilepsy"

### Supplementary Materials #1

**Rationale**: A key hypothesis motivating this study is that connectivity–language relationships may include both (i) **shared effects** that are present in children with and without SeLECTS and (ii) **SeLECTS-related deviations**, in which the connectivity–language association differs in the epilepsy group, potentially reflecting pathological disruption or compensatory mechanisms. However, it is also possible that there are no group-specific associations between connectivity and language. To explicitly test whether modeling group-specific associations provides incremental predictive value, we compared our original MVPA approach with-interaction features to an otherwise identical MVPA pipeline without interaction-features.

**Data**: The same cohort as the main analysis (27 children with SeLECTS and 29 controls), each with both task EEG and resting-state EEG recordings.

**Methods**: We compared two MVPA pipelines that were identical in two-step estimation, nested cross-validation strategy, and evaluation metrics. The only difference was whether interaction features were included in the modeling input.

**Results (Supplementary Figure 1)**: Both models with (task RMSE = 10.88 ± 0.62; rest RMSE = 12.53±1.03) and without interaction features (task RMSE = 12.411 ± 0.646; rest RMSE = 14.174 ± 0.930) outperformed the chance level model (chance level RMSE = 14.779; all p-values $< 0.001$), and models with interaction features outperformed models without interaction features (comparison of task models p-value $< 0.001$; comparison of rest models p-value $= 0.025$). Incorporating interaction features improves out-of-sample prediction.

**Conclusion**: This supplementary comparison supports our modeling choice to include group-modulated interaction features. While primary connectivity features alone predict language above chance, the improved performance of our original method demonstrates that explicitly modeling SeLECTS-related deviations provides additional predictive information.

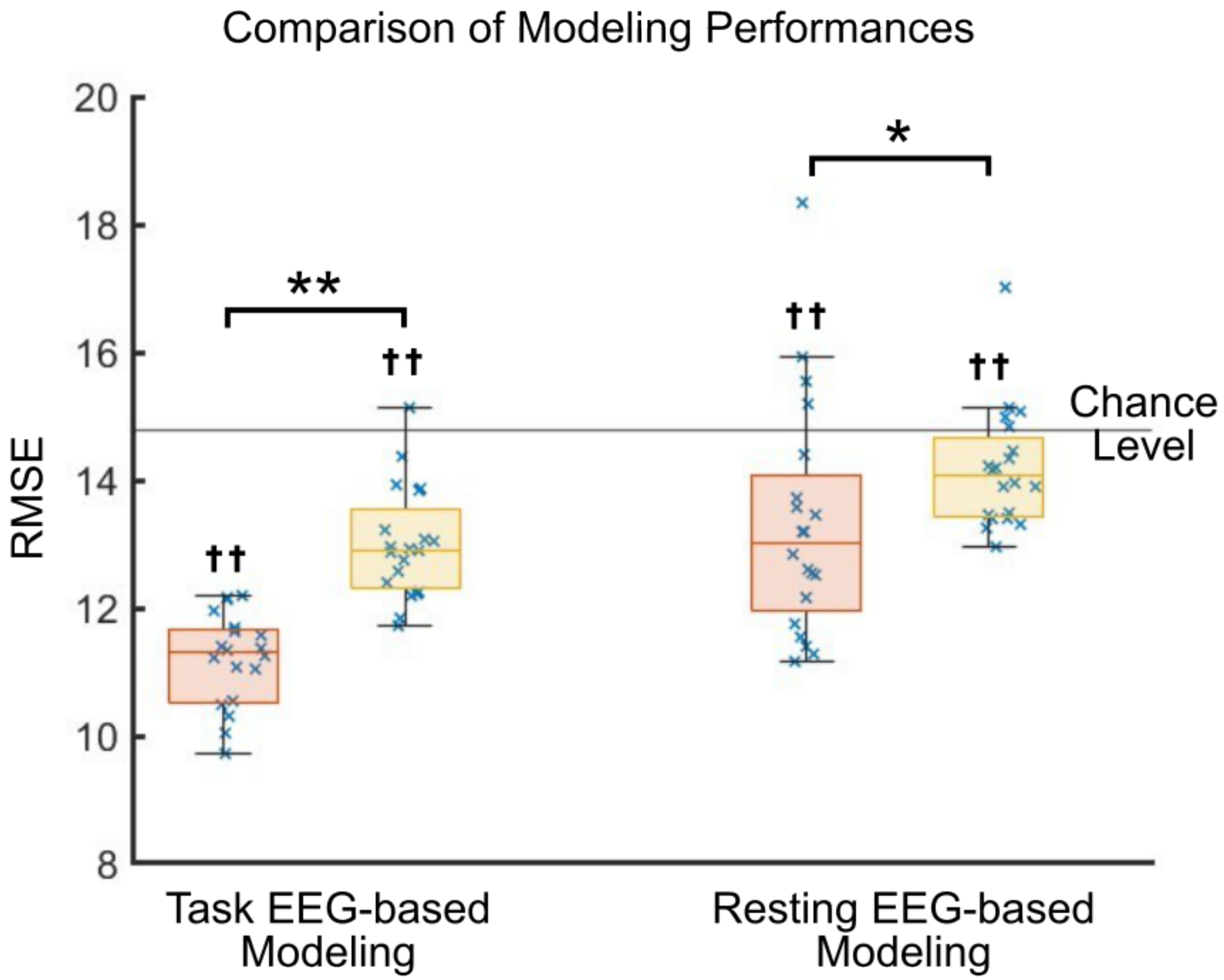


Supplementary Figure 1: MVPA prediction performance for task-based and resting-state EEG connectivity. Each cross represents the RMSE from one repeated nested cross-validation run. Orange boxplots show the RMSE distributions for the original method (with interaction features), and yellow boxplots show the RMSE distributions for the compared connectivity-only method. †† represents $p<0.001$ comparing to the chance level; $**$ represents $p < 0.001$ in the paired comparisons; and $*$ represents $p < 0.05$ in the paired comparisons.

### Supplementary Materials #2

**Rationale**: We describe the weighted MVPA pipeline, including our weighting function, weighted RMSE normalization, and hyperparameter tuning procedure.

**Data**: The same cohort as the main analysis (27 children with SeLECTS and 29 controls), focusing on the task EEG as it predicts language better than the resting EEG.

**Methods**: We implemented a weighted version of the MVPA pipeline described in the main manuscript, using the same input design matrix [$X$, $X .* g$, $g$]. The key modification is introduction of observation weights based on each subject's language score.

Specifically, within each inner cross-validation training fold, each subject is assigned an observation **weight** $w$ according to $w$ = 1+*strengt*h * *max*(0, 100-*y*), where 100 is the population mean of the CTOPP-2 standardization sample (50th percentile normative anchor), whereas $y$ is the actual CTOPP scores of that subject. Participants scoring at or above 100 receive a weight of 1. Participants scoring below 100 are progressively upweighted in proportion to the distance of their score from 100, reflecting the clinical rationale that children performing below the normative mean are more likely to have phonological difficulties with functional/academic consequences. The variable *strengt*h is a hyperparameter controlling the observation weight $w$.

*Weighted RMSE for model evaluation*

The *standard RMSE* weighs all prediction errors equally across subjects. A model judged by the standard RMSE can therefore achieve good overall performance while systematically underperforming for children at the lower end of the score distribution. To evaluate model performance specifically in lower-scoring children, we compute a *weighted RMSE*, in which each subject's squared prediction error is multiplied by their observation weight before averaging.

Notably, both the primary MVPA model (presented in our main manuscript) and the weighted MVPA model (presented in this supplementary analysis) produce predicted language scores for each participant. As such, we can calculate both *standard RMSE* and *weighted RMSE* values for both models, allowing us to directly and fairly compare the primary and weighted models to each other. Weighted and overall RMSE are both reported to quantify the trade-off between optimizing for the full distribution vs. lower-scoring children.

*Hyperparameter tuning*

Since changing the hyperparameter *strengt*h changes the overall scale of the weights (and hence the scale of the weighted error metrics), we cannot use the weighted RMSE to tune the strength hyperparameter. Instead, we tuned this hyperparameter using the normalized weight RMSE. Specifically, for each candidate strength, the raw weights were normalized within each fold by dividing by their mean so that the average weight equals 1. These normalized weights were applied consistently in the RFE feature-selection criterion, the ridge regression loss function, and the inner cross-validation evaluation used for hyperparameter tuning. This normalization ensures that normalized weighted RMSE values can be served as a concrete criterion and are directly comparable across candidate strength values.

The *strengt*h was tuned within the inner cross-validation loop via a grid search over the range 0.1 to 1.0 (step size 0.1), with additional candidate values of 0.01, 0.05, 2, 4, 6, and 10. For each candidate strength, we (i) computed and normalized observation weights using only the inner-training data, (ii) performed the full two-step estimation procedure (RFE and ridge) with those weights, and (iii) evaluated performance on inner-validation folds using normalized weighted RMSE as the selection criterion.

**Results**: The value *strengt*h = 1 was selected given the lowest average normalized weighted RMSE across inner-validation folds (Supplementary Figure 2).

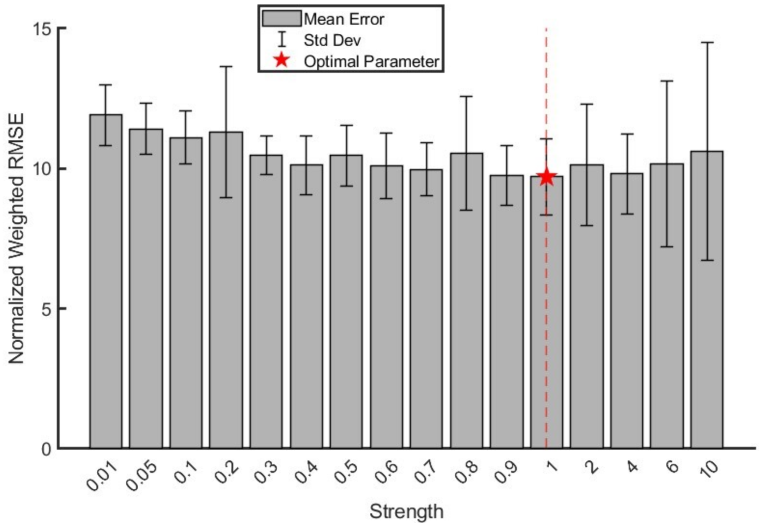


Supplementary Figure 2: Hyperparameter selection based on normalized weighted RMSE (nwRMSE). Bar plots show the mean nwRMSE for each of the candidate hyperparameter settings, calculated across 20 repeated model runs. Error bars represent the standard deviation of the nwRMSE across repetitions (mean ± SD). The optimal hyperparameter was identified as the minimum average nwRMSE.

### Supplementary Table and Figures:

Supplementary Table 1: Data quality and retained trials after preprocessing for children with SeLECTS and controls. P-values are from Wilcoxon signed-rank tests compared between groups (SeLECTS vs. Controls).

| Total (n=56) | SeLECTS (n=27) | Controls (n=29) | p-value |
|---|---|---|---|
| Trials Retained (mean ± SD) | 47.5±6.3 | 49.3±2.0 | 0.165 |
| Rejected EEG Channels (mean ± SD) | 5.6 ±2.0 | 5.4±2.2 | 0.721 |
| Rejected Independent Components (mean ± SD%) | 23.4±11.1% | 20.4±5.2% | 0.111 |
| Verb Accuracy (mean ± SD) | 0.785±0.162 | 0.270±0.187 | 0.196 |

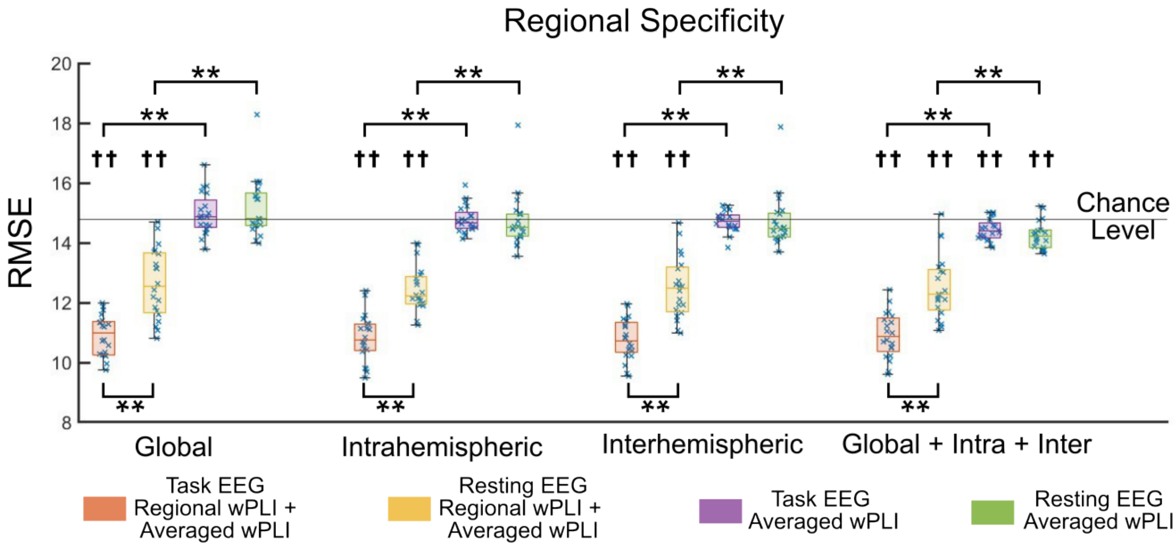


Supplementary Figure 3: Prediction performance, measured by root mean squared error (RMSE), of the tested models. Box plots depict the distribution of these performances. Orange boxes: model inputs as regional connectivity calculated from task EEG combined with the averaged measures (i.e., global; intrahemispheric; interhemispheric averaged connectivity); Yellow boxes: model inputs as regional connectivity calculated from resting EEG combined with the averaged measures; purple boxes: model input was the averaged connectivity measures from task EEG alone; green boxes: model input was the averaged connectivity measures from resting EEG alone. †† represents significant after multiple comparison correction comparing to the chance level; and ** represents significant after multiple comparison correction in the paired comparisons. ‡‡ represents significant after multiple comparison correction in the paired comparison between task and resting EEG based connectivity.

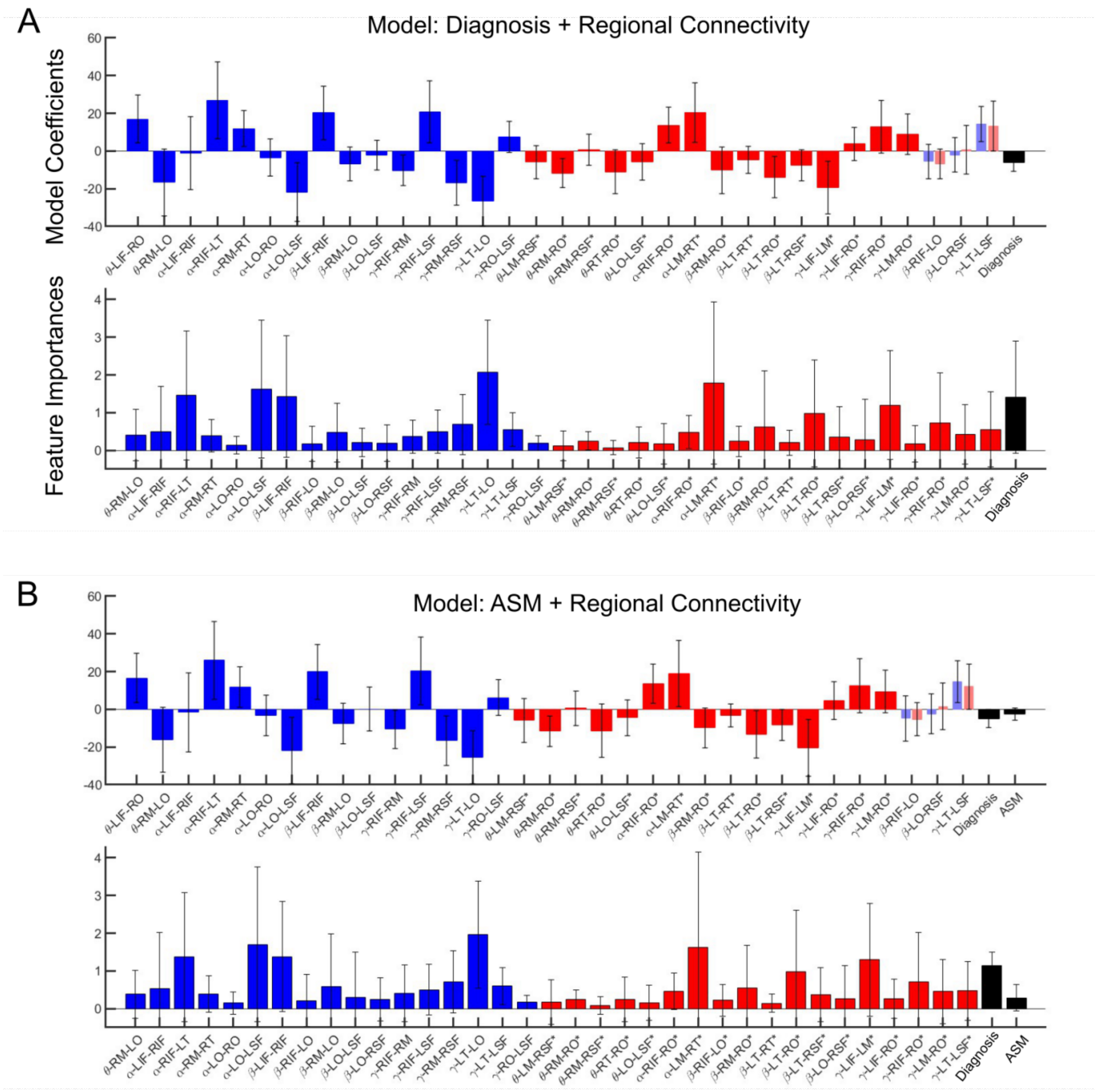


Supplementary Figure 4: Model coefficients and PFI for task-EEG connectivity predictors. A: Language-prediction model using the interaction-augmented regional connectivity design with diagnosis/group indicator. B: Language-prediction model using the same interaction-augmented regional connectivity design with antiseizure medication (ASM). For each panel, the **top subplot** shows model coefficients for the retained predictors: blue bars denote the primary (control-referenced) coefficients $\beta_j$, red bars denote the deviation (SeLECTS–control) coefficients $\gamma_j$. The **bottom subplot** shows permutation feature importance. Blue bars indicate predictors interpreted as shared patterns, whereas red bars indicate predictors interpreted as SeLECTS-specific patterns. Black bars indicate the PFI of the clinical covariate(s) included in the design (group/diagnosis in A; ASM in B).

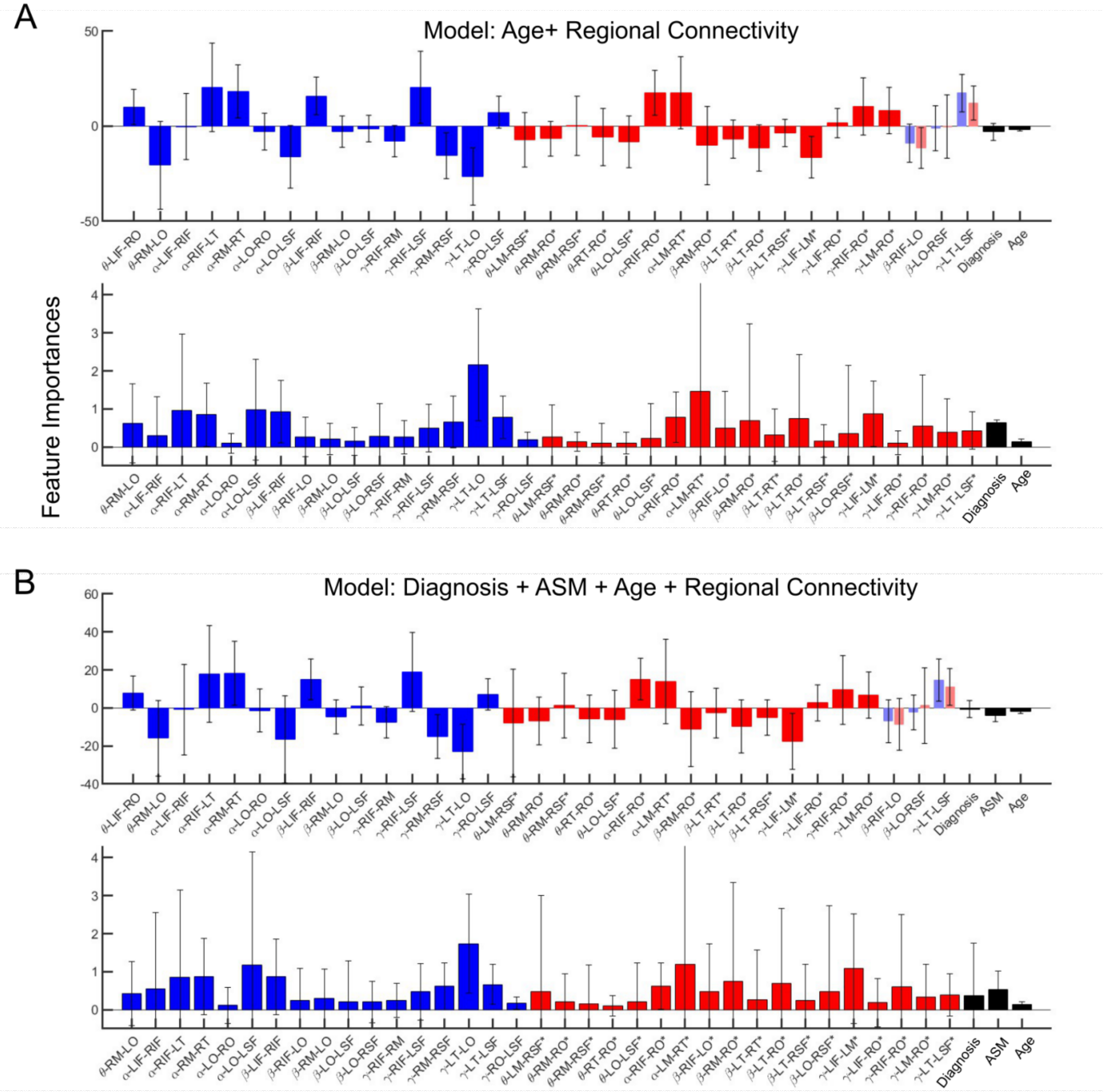


Supplementary Figure 5: Model coefficients and PFI for task-EEG connectivity predictors. A: Language-prediction model using the interaction-augmented regional connectivity design with diagnosis/group indicator. B: Language-prediction model using the same interaction-augmented regional connectivity design with antiseizure medication (ASM). For each panel, the **top subplot** shows model coefficients for the retained predictors: blue bars denote the primary (control-referenced) coefficients $\beta_j$, red bars denote the deviation (SeLECTS–control) coefficients $\gamma_j$. The **bottom subplot** shows PFI. Blue bars indicate predictors interpreted as shared patterns, whereas red bars indicate predictors interpreted as SeLECTS-deviating patterns. Black bars indicate the PFI of the clinical covariate(s) included in the design (Age in A; [Diagnosis, ASM, Age] in B).

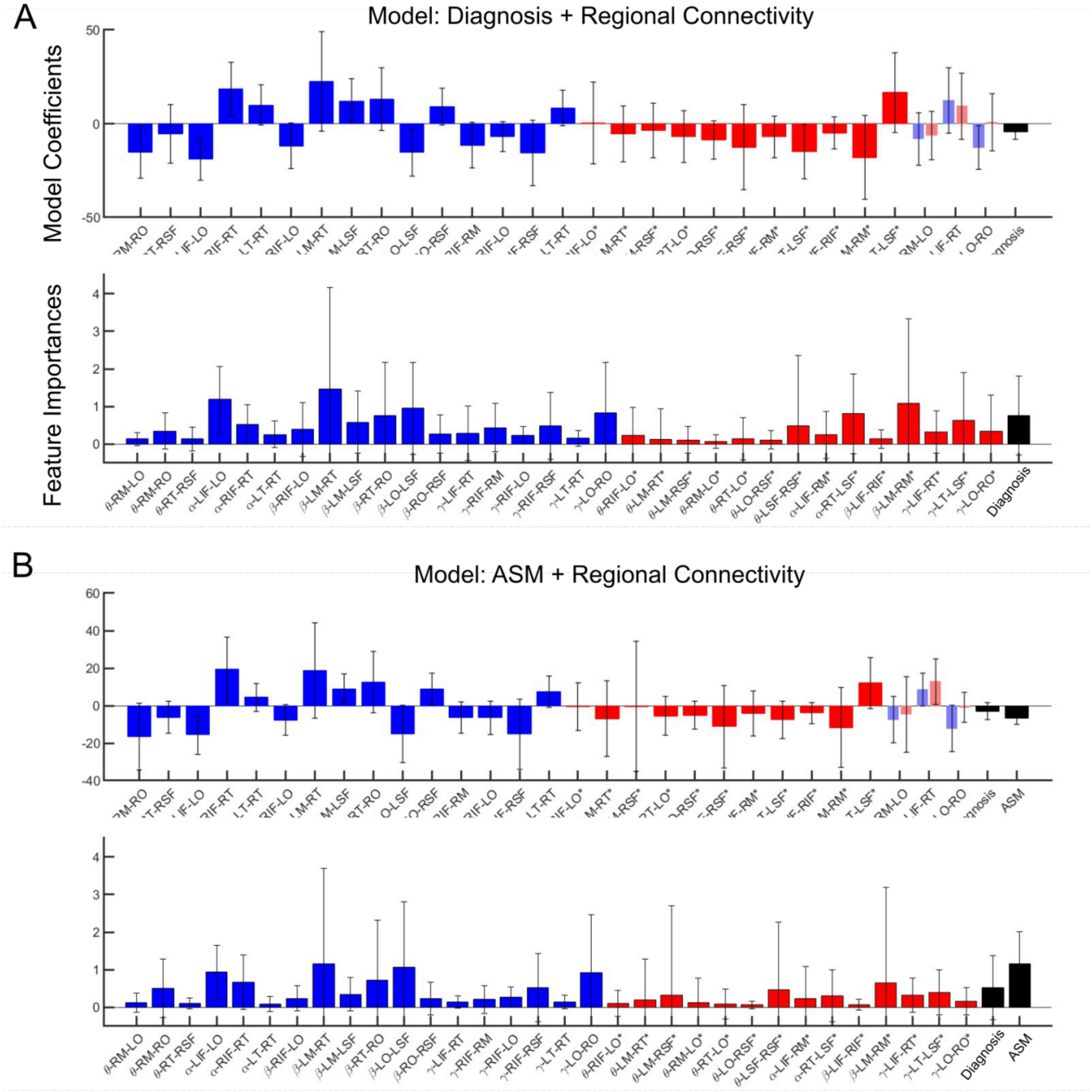


Supplementary Figure 6: Model coefficients and PFI for resting-EEG connectivity predictors. A: Language-prediction model using the interaction-augmented regional connectivity design with diagnosis/group indicator. B: Language-prediction model using the same interaction-augmented regional connectivity design with antiseizure medication (ASM). For each panel, the **top subplot** shows model coefficients for the retained predictors: blue bars denote the primary (control-referenced) coefficients $\beta_j$, red bars denote the deviation (SeLECTS–control) coefficients $\gamma_j$. The **bottom subplot** shows permutation feature importance. Blue bars indicate predictors interpreted as shared patterns, whereas red bars indicate predictors interpreted as SeLECTS-deviating patterns. Black bars indicate the PFI of the clinical covariate(s) included in the design (group/diagnosis in A; ASM in B).

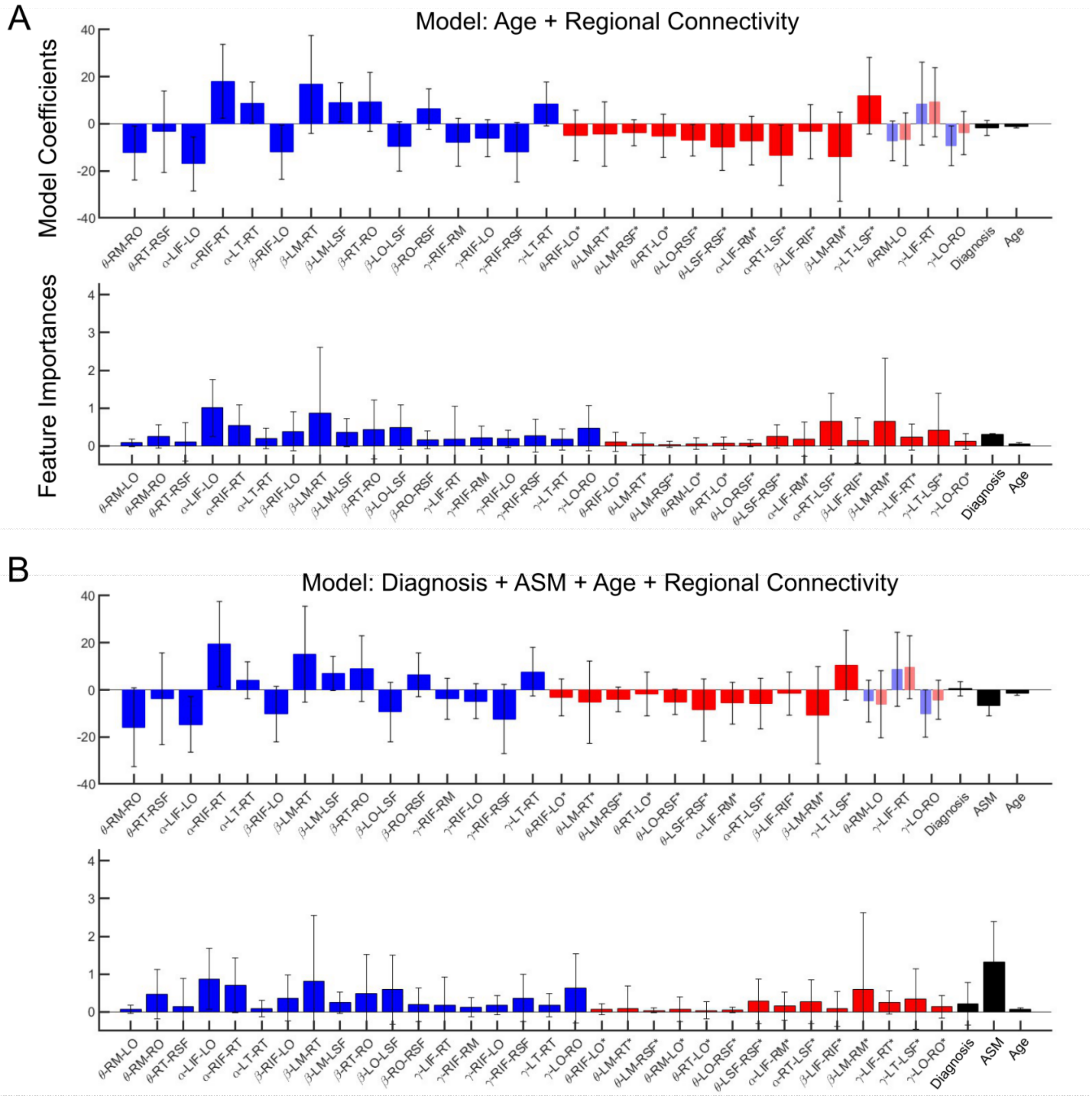


Supplementary Figure 7: Model coefficients and PFI for resting-EEG connectivity predictors. A: Language-prediction model using the interaction-augmented regional connectivity design with diagnosis/group indicator. B: Language-prediction model using the same interaction-augmented regional connectivity design with antiseizure medication (ASM). For each panel, the **top subplot** shows model coefficients for the retained predictors: blue bars denote the primary (control-referenced) coefficients $\beta_j$, red bars denote the deviation (SeLECTS–control) coefficients $\gamma_j$. The **bottom subplot** shows PFI. Blue bars indicate predictors interpreted as shared patterns, whereas red bars indicate predictors interpreted as SeLECTS-deviating patterns. Black bars indicate the PFI of the clinical covariate(s) included in the design (Age in A; [Diagnosis, ASM, Age] in B).
